# Uncovering disease-related multicellular pathway modules on large-scale single-cell transcriptomes with scPAFA

**DOI:** 10.1101/2024.03.11.584023

**Authors:** Zhuoli Huang, Yuhui Zheng, Weikai Wang, Wenwen Zhou, Chen Wei, Xiuqing Zhang, Xin Jin, Jianhua Yin

## Abstract

Pathway analysis is a crucial analytical phase in disease research on single-cell RNA sequencing (scRNA-seq) data, offering biological interpretations based on prior knowledge. However, currently available tools for generating cell-level pathway activity scores (PAS) exhibit computational inefficacy in large-scale scRNA-seq datasets. Besides, disease-related pathways are commonly identified by cross-condition comparisons in each cell type, neglecting the potential multicellular patterns. Here, we present single-cell pathway activity factor analysis (scPAFA), a Python library designed for large-scale single-cell dataset allowing rapid PAS computation and uncovering biologically interpretable disease-related multicellular pathway modules, which are low-dimensional representations of disease-related PAS variance in multiple cell types. Application on colorectal cancer (CRC) dataset with 371,223 cells and large-scale lupus atlas over 1.2 million cells demonstrated that scPAFA can achieve > 33-fold decreases in runtime of PAS computation and further identified reliable and interpretable multicellular pathway modules that capture the transcriptomic features of CRC tumor status and transcriptional abnormalities in lupus patients, respectively.

---

Single-cell RNA sequencing (scRNA-seq) technologies have revolutionized molecular biology by facilitating the high-throughput and high-resolution profiling of single-cell level transcriptomes^1^. In the past few years, the prevalence of scRNA-seq and the implementation of sample multiplexing technique^2^ have led to the emergence of large-scale single cell transcriptome atlas. Representative examples of large-scale atlases containing peripheral blood mononuclear cell(PBMC) atlas with over 1.2 million cells from 162 systemic lupus erythematosus(SLE) cases and 99 healthy controls^3^; COVID-19 comprehensive immune landscape with 1.46 million cells from 196 individuals^4^; integrated cell atlas of the lung in health and disease spanning over 2.4 million cells from 486 individuals^5^; adult human brain dataset comprised more than 3 million cells, including more than 2 million neurons^6^. These atlases function as indispensable resources for delving into the intricacies of both healthy and diseased cellular states; nevertheless, they concurrently place higher requirements on the stability and efficiency of analytical methods. Pathway analysis constitutes a pivotal analytical phase in the interpretation of omics data, facilitating the detecting of alterations in cellular biological processes^7^. Existing single-cell pathway activity scoring methods, such as AddModuleScore^8, 9^, UCell^10^, and AUCell^11^ can generate pathway activity scores (PAS) for individual cells, rather than cell populations or clusters, and these scores can then be used for downstream analysis^7^, such as clustering, cell type identification and inter-group comparisons based on biological conditions (e.g., case-control comparisons). Most single-cell studies analyze pathway activation based on curated gene lists compiled by domain experts, which represent the current reference biological knowledge^8^. In contrast to utilizing the expression levels of individual genes, pooling of gene set based measurements, which amalgamate the functional effects of diverse genes participating in identical biological pathways, can significantly augment statistical robustness and facilitate biological interpretation when discerning specific cellular functions or states, especially in sparse and noisy scRNA-seq dataset^12, 13, 14^. Nonetheless, abovementioned methods exhibit tardiness and computational inefficacy in large-scale scRNA-seq datasets, especially in situations where a large number of pathways is required to be calculated.

A typical primary step in the analysis of scRNA-seq data is to partition the cells into clusters and annotate as cell types^15^; therefore, downstream analysis commonly focuses on cell-type-centric pairwise cross-condition comparisons, disregarding the multicellular nature of disease processes^16^.

We assumed that aggregating cross-condition differences of pathway activity scores detected in various cell types into a low-dimensional module, would enable a more comprehensive elucidation and interpretation of the transcriptomic features representing the disease state. It was reported that multi-omics factor analysis (MOFA) is a statistical framework for the comprehensive and scalable integration of multi-modal data, which can analyze data from complex experimental designs that include multiple data modalities and multiple groups of samples, such as batch or experimental conditions^17, 18^. Gaining advantages from the high flexibility of statistical framework, MOFA was widely applied in different biological contexts, for instance, revealing an axis of heterogeneity associated with the disease outcome in chronic lymphocytic leukemia^19^; identification of alterations in Alzheimer’s disease^20^; uncovering genotypic, microbiome, and metabolomic factors associated with adenoma and colorectal cancer risk^21^. Inspired by Ramirez et al^16^, we reutilized MOFA to identify multicellular pathway modules. Here, we present single-cell pathway activity factor analysis (scPAFA), an open-source python library designed for large-scale single-cell dataset to rapidly compute PAS and uncover disease-related MOFA-based multicellular pathway modules. The scPAFA contains time efficient implementations of single-cell pathway activity scoring algorithms, which can compute PAS for over 1300 pathways in million-cell-level scRNA-seq data within 40 minutes. It also provides user-friendly functions for PAS matrices preprocessing according to different experimental designs, MOFA model training and downstream analysis of multicellular pathway modules. As a case study, we benchmarked scPAFA’s performance on a colorectal cancer (CRC) dataset^22^ together with a large-scale lupus atlas^3^. We have demonstrated that scPAFA found trustworthy and interpretable multicellular pathway modules which represent the transcriptomic characteristic of CRC tumors status and transcriptional abnormalities in patients with SLE, respectively. Furthermore, high-weight features in the modules demonstrate outstanding performance in machine learning as input for classifier training. All documentation, tutorials and source code for scPAFA can be found at https://github.com/ZhuoliHuang/scPAFA.

## RESULTS

### Overview of scPAFA workflow

The scPAFA workflow consists of four steps (Fig. 1A and Methods), each step of scPAFA workflow is supported by user-friendly application programming interface (API) allowing customized parameters. In the first step, single-cell gene expression matrix and collection of pathways are used to compute PAS by ‘fast_ucell’ or ‘fast_score_genes’. These functions are more computationally efficient implementation of UCell and AddModuleScore (also known as ‘score_genes’ in Scanpy^23^), which are commonly used single-cell pathway activity scoring methods. Collection of pathways can be obtained from public databases, such as the Molecular Signatures Database (MsigDB)^24, 25^ and National Center for Advancing Translational Sciences (NCATS) BioPlanet^26^, and customized based on specific biological context of interest. Second, single-cell PAS matrix is reformatted into a suitable input for MOFA along with cell-level metadata including sample/donor, cell type and technical batch information. In brief, we reutilized multi-group MOFA+ framework by assigning pathways as features, cell types as non-overlapping sets of modalities (also called views) and technical batch information as groups. Features will be centered per group before fitting the model to reveal which sources of variability are shared between the different groups, therefore batch effects are mitigated. In each group and view, cell-level PAS is aggregated into pseudobulk-level PAS by samples/donors. In the third step, MOFA+ model is trained and the integrated latent factors matrix associated with feature weights matrix that explain the major axes of variation in PAS across the datasets is extracted from the converged model. Finally, along with the sample-level clinical metadata, disease-related multicellular pathway modules (latent factor and its corresponding weights of pathways across cell types) can be identified by statistical analysis. Downstream analyses include characterizing and interpreting of multicellular pathway modules, samples/donors stratification, classifier training based on high-weight pathways.

**Figure 1.**
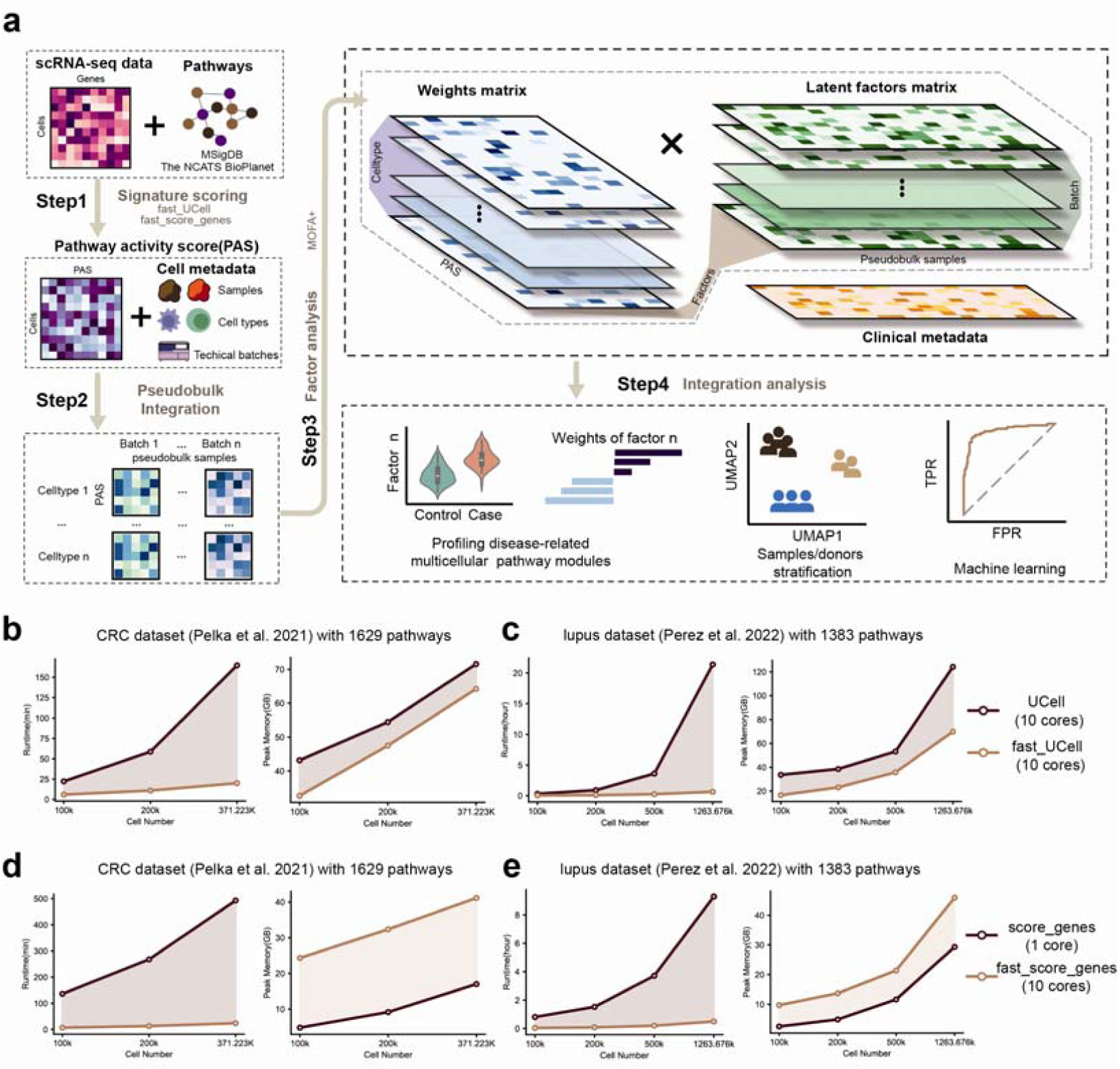
scPAFA overview and its performance on CRC and lupus dataset. (a) Schematic outline of scPAFA workflow, which consists of four steps: 1) Generating PAS matrix from single cell gene expression matrix and pathway set using ‘fast_ucell’ or ‘fast_score_genes’ functions. 2) Aggregating cell-level PAS matrix into pseudobulk-level PAS long data integrating with sample, cell type and batch information. 3) Training MOFA model based on experimental design 4) Using latent factor matrix and its corresponding weight matrix of MOFA model to identify disease related multicellular pathway modules and provide biological interpretation. (b) Line graph shows the runtime and memory usage of UCell and ‘fast_ucell’(scPAFA) on CRC scRNA-seq dataset with 1629 pathways. (c) Line graph shows the runtime and memory usage of UCell and ‘fast_ucell’(scPAFA) on lupus scRNA-seq dataset with 1383 pathways. (d) Line graph shows the runtime and memory usage of ‘score_genes’(Scanpy) and ‘fast_score_genes’(scPAFA) on CRC scRNA-seq dataset with 1629 pathways. (e) Line graph shows the runtime and memory usage of ‘score_genes’(Scanpy) and ‘fast_score_genes’(scPAFA) on lupus scRNA-seq dataset with 1383 pathways.

### Validating scPAFA using scRNA-seq data

To show scPAFA’s performance, we benchmarked it on two public scRNA-seq datasets in different diseases: colorectal cancer (CRC) dataset with 371,223 cells, collected from colorectal tumors and adjacent normal tissues of 28 mismatch repair-proficient (MMRp) and 34 mismatch repair-deficient (MMRd) individuals^22^; lupus dataset with 1,263,676 cells, collected from PBMCs of 162 SLE cases and 99 healthy controls^3^. We used NCATS BioPlanet^26^, a single collection of known biological pathways operating in human cells which incorporates 1,658 pathways as input. Specifically, for the CRC dataset, we additionally included 149 gene sets mined from the Curated Cancer Cell Atlas (3CA) metaprogram^27^. After quality control, 1629 and 1383 pathways were taken as input for CRC dataset and lupus dataset, respectively. We first examined the PAS computation speed of scPAFA on an Intel X79 Linux server using 10 cores. It was observed that the running speeds of functions ‘fast_ucell’ and ‘fast_score_genes’ were approximately 3.8 to 33.1 times faster than their original versions on dataset range from 10,000 cells to 1,263,676 cells (Fig. 1b-e). For instance, on complete lupus dataset, UCell cost 21.4 hours for 1383 pathways, whereas ‘fast_ucell’ cost 38.73 minutes(Fig. 1c). Compared to existing methods, scPAFA can achieve greater computational efficiency gains on large-scale data set for pathway activity scoring. In addition, the peak memory usage of ‘fast_ucell’ was lower than UCell(Fig. 1b, c). However, the memory usage of ‘fast_score_genes’ was higher than ‘score_genes’, due to memory overhead of concurrent execution, whereas original ‘score_genes’ function can only utilize a single core(Fig. 1d, e).

### Multicellular pathway modules in tumor cells that stratify CRC patients

CRC is one of the most common cancers in developed countries. Immune responses to CRC are highly variable, tumors with a mismatch repair-deficiency (MMRd)/microsatellite instability-high (MSI-H) phenotype shows unique characteristics, including elevated tumour mutational burden and enhanced anti-tumor immunity than mismatch repair-proficient (MMRp) tumors^22, 28^. Based on single cell data and cell type annotations from abovementioned CRC dataset^22^, we further used scPAFA to identify MMR status related multicellular pathway modules. In brief, we extracted the 108,497 epithelial tumor cells from the CRC dataset, which were reported as 11 cell clusters. The PAS matrix with 1629 pathways was computed by ’fast_ucell’, then aggregated along with samples and cell types information. We first used single-group MOFA framework and revealed obvious batch effect in samples’ stratification caused by different 10X chemistry version, which was corresponding to the protocol for generating single cell gene expression libraries (Supplementary Fig. 1a, b); hence PAS matrix was reformatted with batch, samples and cell type information while multi-group MOFA+ framework was further applied. The converged model provided a latent factor matrix with 8 factors that explain at least 1% of variance and 65 pseudobulk-samples, the Uniform Manifold Approximation and Projection (UMAP) were used to perform dimensionality reduction and visualization on the latent factor matrix, showing distinct stratification between MMRd and MMRp CRC patients. The values of factor2 and factor3 were significantly higher in MMRd samples, whereas factor6 demonstrated opposite trends (Fig .2a). Noteworthy, pseudobulk-samples from different batches exhibited non-segregated distribution on the UMAP plot, and the factor values showed no significant differences between batches, suggesting that no batch-related factors were detected (Fig .2b). Moreover, two-dimensional scatter plot based on the combination of factors 2, 3, and 6 also showed stratification of pseudobulk-samples by MMR status (Fig .2c). Simultaneously, hierarchical clustering based on the values of factors 2, 3, and 6 also revealed MMR status related stratification, additionally, heterogeneity among samples in each MMR status was also detected (Fig .2d). The variance captured by factors2, 3 and 6 was mainly contributed by stemTA-like cell clusters (cE01, cE02 and cE03) and immature goblet (cE06) (Fig .2e). Besides, factor2 positively correlates with factor3, while factor6 negatively correlates with both factor2 and factor3. In conclusion, the above results suggest that factors 2, 3 and 6 can be considered as tumor cell-based MMR status related multicellular pathway modules.

**Figure 2.**
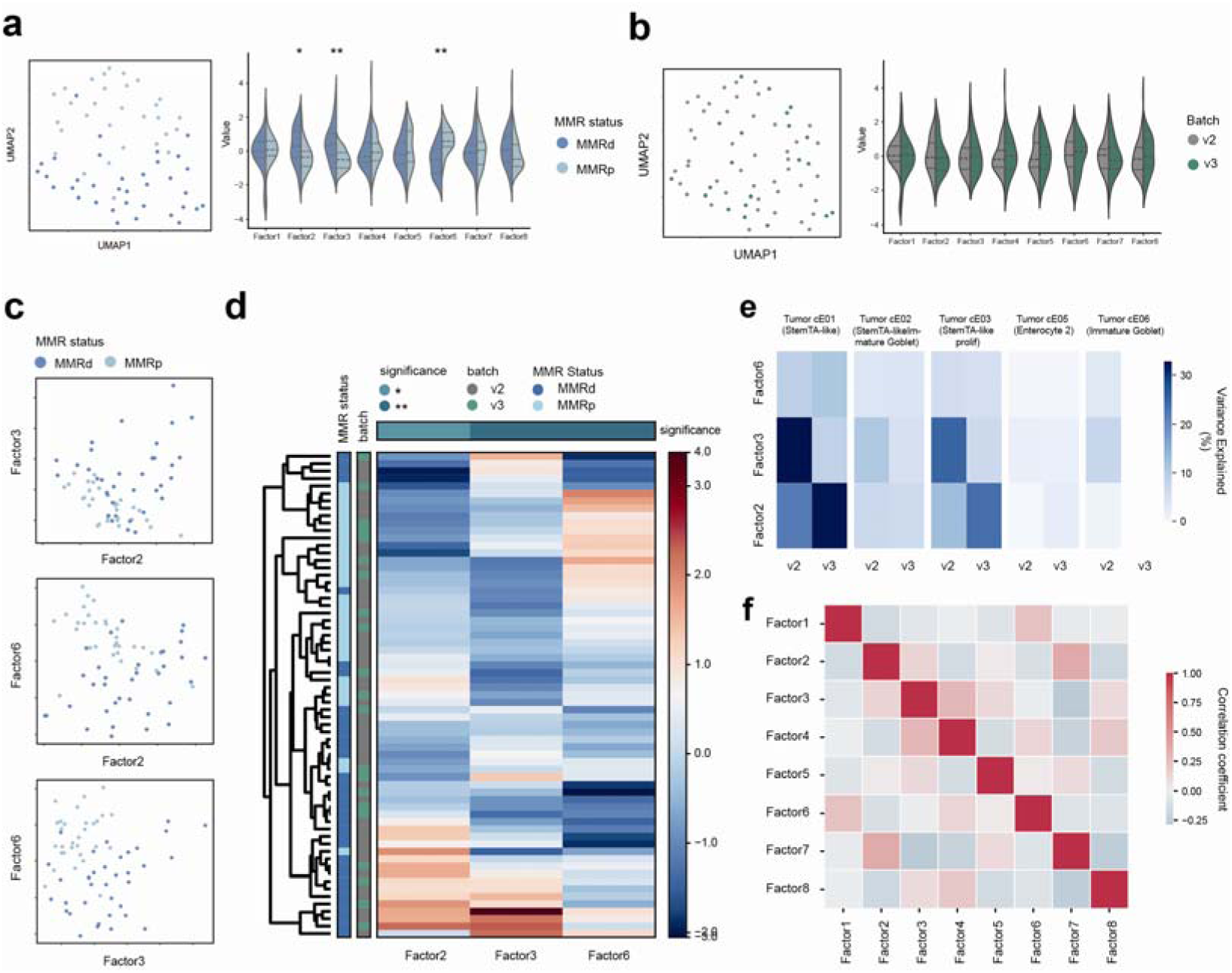
Multicellular pathway modules stratify CRC patients. (a) UMAP of latent factor matrix shows stratification between MMRp and MMRd samples (left). Violin plots (right) shows the difference of factor values between MMRp and MMRd samples using Mann-Whitney U test. * adjust p-value < 0.05, **adjust p-value < 0.01. (b) UMAP of latent factor matrix shows uniform distribution of samples from different batches (left). Violin plots (right) shows the difference of factor values between batch v2 and batch v3 samples using Mann-Whitney U test. * adjust p-value < 0.05, **adjust p-value < 0.01. (c) Scatter plots of factor 2, 3 and 6 show stratification between MMRp and MMRd samples. (d) Heatmap with hierarchical clustering of factor 2, 3 and 6 shows stratification between MMRp and MMRd samples. The annotation of columns (top) shows the difference of factor values between MMRp and MMRd samples using Mann-Whitney U test. * adjust p-value < 0.05, **adjust p-value < 0.01. (e) The heatmap displays percentage of variance explained for each factor across the different groups (batch) and views (cell type). For simplicity, only the factor 2, 3 and 6 was shown. (f) The heatmap displays correlation coefficient between different factors.

### Pathway-centric interpretation of MMR status related modules

We further employed the feature weights matrixes corresponding to the factors 2, 3 and 6 to interpret MMR status related multicellular pathway modules. Weights provide a score indicating the strength of the relationship between each feature and factor, for instance, pathway-cell type pairs with no association with the factor have values close to zero, while pathway-cell type pairs with stronger association have larger absolute values. The sign of the weight indicates the direction of the effect, consistent with the direction of the factor values. MMRd related multicellular pathway modules consists of ‘eosinophils in the chemokine network of allergy’ in immature goblet, ‘METAPROGRAM_17_INTERFERON_MHC_II_1’ in stemTA-like immature goblet and ‘METAPROGRAM_6_HYPOXIA’ in stemTA-like immature goblet, etc., whereas MMRp related multicellular pathway modules included ‘METAPROGRAM_B_CELLS_METABOLISM_MYC’ in stemTA-like cells, ‘Cap-dependent translation initiation’ in enterocyte 2 and ‘Nef-mediated downregulation of MHC class I complex cell surface expression’ in stemTA-like immature goblet, etc. (Fig. 3a, Supplementary Table 1) Furthermore, we validate the PAS differences between MMRd and MMRp of some high-weight pathway-cell type pairs extracted from feature weights matrixes corresponding to the factors 2,3 and 6. We revealed that ‘METAPROGRAM_17_INTERFERON_MHC_II_1’, a high-weight pathway associated with factors 2 and 3, PAS of which was significantly increased in MMRd patients across various cell types comparing with MMRp patients (Fig. 3b). Pelka et al^28^ reported that ISG and MHC class II gene programs were more active in MMRd versus MMRp tumors, which was consistent with our results. It is widely reported that MMRd CRC patients are more sensitive in response to anti-programmed death-1 receptor (PD-1)/programmed death-1 receptor ligand 1 (PD-L1) therapy than MMRp patients^29, 30, 31^, consistently, PAS of ‘PD-1 signaling’ pathway in immature goblet that associated with factor 2 was significantly elevated in MMRd patients (Fig. 3b). Moreover, PAS of factor 6 associated-pathway ‘METAPROGRAM_6_HYPOXIA’ was significantly up-regulated in MMRd patients, which was consistent with previous research reporting hypoxia causes downregulation of mismatch repair system in cells^32, 33, 34^(Fig. 3b). In addition, we also revealed that significant PAS up-regulation of factor 6 associated-pathway ‘Glycolysis’ in MMRd patients, similar to the results obtained by Pelka et al^22^. Overall, pathway-centric interpretation of MMR status related multicellular modules substantiated their validity and trustworthiness.

**Figure 3.**
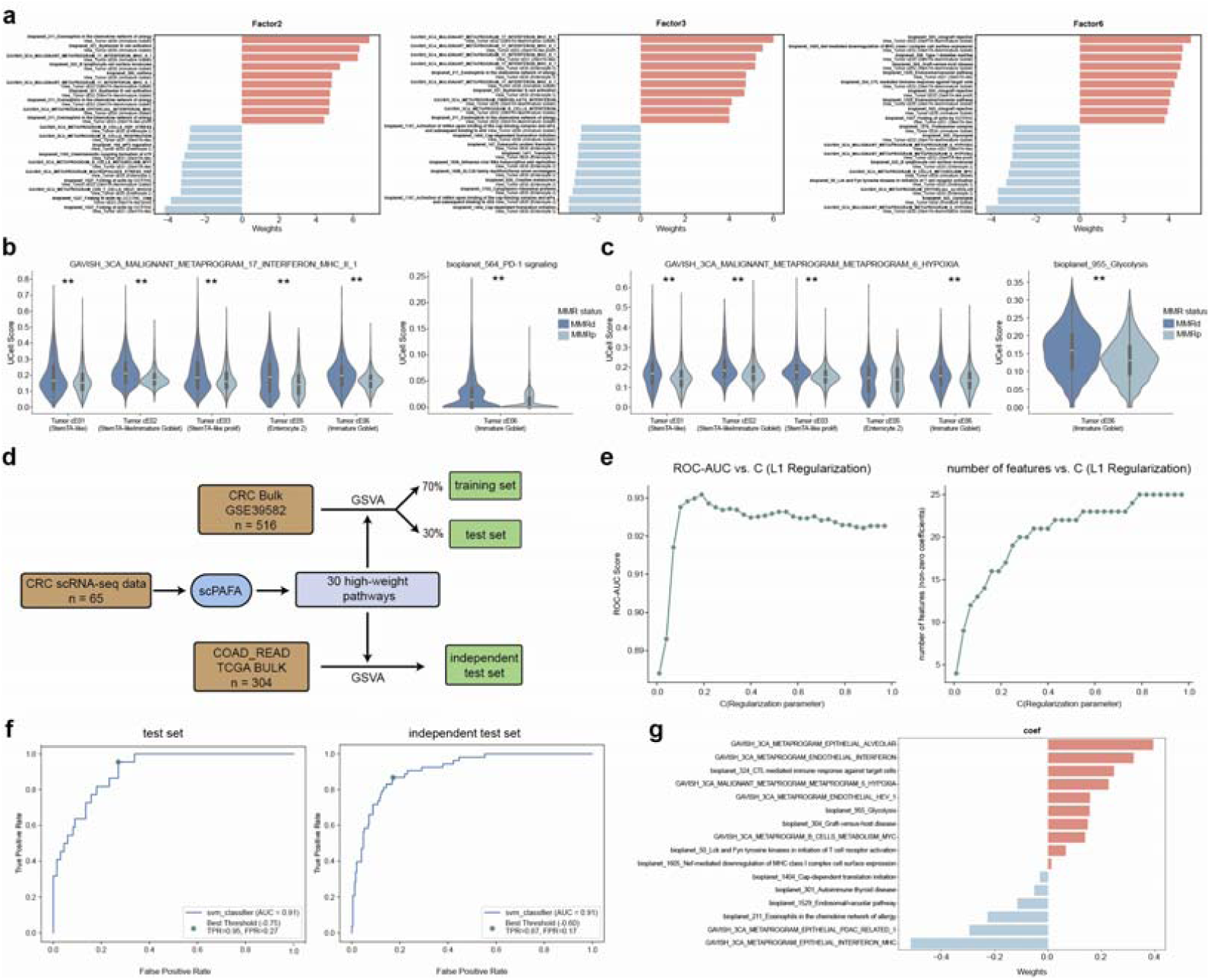
Pathway-centric interpretation and classifier training based on MMR status related modules. (a) Butterfly bar plots displaying the pathway-cell type pairs with top 10 positive and negative weights of factor 2, 3 and 6. (b) Violin plots show the difference of cell-level PAS of METAPROGRAM_17_INTERFERON_MHC_II_1’(left) and ‘PD-1 signaling’(right) between MMRp and MMRd using Mann-Whitney U test. * p-value < 0.05, ** p-value < 0.01. (c) Violin plots show the difference of cell-level PAS of ‘METAPROGRAM_6_HYPOXIA’(left) and Glycolysis’(right) between MMRp and MMRd using Mann-Whitney U test. * p-value < 0.05, ** p-value < 0.01. (d) Flowchart for classifier training on bulk RNA data. (e) Line chart (left) illustrates the impact of various regularization parameter values on AUROC in the training set (4-fold cross-validation). The line chart (right) illustrates the impact of different regularization parameter values on the number of features in the training set (4-fold cross-validation). (f) The AUROC of the classifier on the test set and an independent test set. (g) Butterfly bar plots display the coefficients of features included in the classifier.

To test the robustness and generality of the multicellular pathway modules uncovered by scPAFA, we used CRC bulk RNA dataset to constructing MMR status classifier based on the high-weight pathways associated with the modules. We applied support vector classification (SVC) model with L1 regularization, using gene set variation analysis (GSVA) score of high-weight pathways as features (Fig. 3d). The L1 regularization parameter was set as 0.2 based on the area under the receiver operating characteristic curve (AUROC) from 4-fold cross-validation in training set (Fig. 3e). The AUROC of MMR status classifier achieved 0.91 in test set (GSE39582^35^) and independent test set (TCGA COAD_READ), besides, the features remained after L1 regularization included pathways such as ‘METAPROGRAM_EPITHELIAL_INTERFERON_MHC’, ‘METAPROGRAM_6_HYPOXIA’ and ‘Glycolysis’ (Fig. 3f). The above results showed the remarkable ability of scPAFA in feature selection.

### Identification of SLE related multicellular pathway modules

SLE is a heterogeneous autoimmune disease, pervious bulk transcriptomic profiling reported that increased type 1 interferon signaling, dysregulated lymphocyte activation, and failure of apoptotic clearance as hallmarks of SLE^3^. For large-scale lupus scRNA-seq atlas, we extracted 1,067,499 cells from 3 processing batches that simultaneously contain SLE and health controls to Identify SLE related multicellular pathway modules by scPAFA workflow. Single-group MOFA framework discovered batch effects in samples’ stratification caused by processing batches (Supplementary Fig. 1c). It’s noteworthy that a cluster (cluster 0) of pseudobulk-samples from batch 3 separated from other pseudobulk-samples, which was characterized by multicellular pathway modules associated with red blood cells (Supplementary Fig. 1c-f). Considering that the lupus dataset was collected from PBMCs, we speculate that cells originating from this cluster have been contaminated; therefore, we excluded these cells, leaving 941,542 cells remaining. The remained dataset was then split into a training set and a test set with a 7:3 ratio based on pseudobulk-samples. For training set, PAS matrix was reformatted with 3 processing batches, 175 pseudobulk-samples and 7 cell types as input for multi-group MOFA+ framework. The UMAP plot of latent factor matrix with 8 factors that explain at least 1% of variance showed distinct stratification between SLE patients and healthy control, while pseudobulk-samples from different batches exhibit a uniform distribution (Fig. 4a, b). Moreover, the value of factor 1 was significantly increased in SLE patients, while factor 6 was significantly decreased (Fig. 4a), hence factors 1 and 6 were identified as SLE related multicellular pathway modules. Consistently, obvious stratification between SLE patients and healthy control can also be observed in two-dimensional scatter plot and hierarchical clustering heatmap based on the values of factors 1 and 6 (Fig. 4c, d). Factor 1 captured variance across all cell types, particularly in classical and non-classical monocytes (cM, ncM), whereas variance captured by factor 6 was mainly contributed by CD4+ T cells and CD8+ T cells (T4, T8) (Fig. 4e). Besides, factor 1 negatively correlates with factor 6 (Fig. 4f).

**Figure 4.**
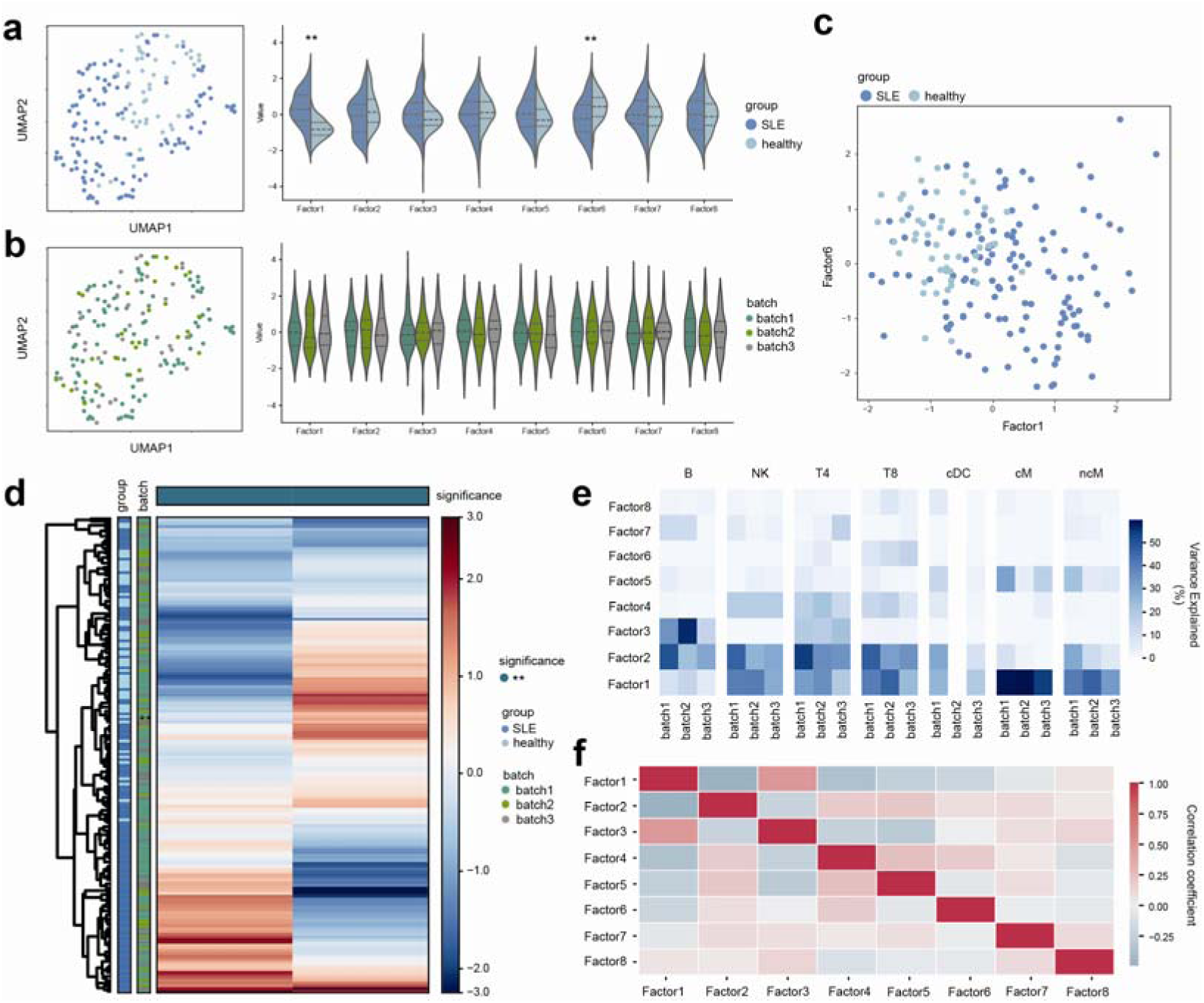
Multicellular pathway modules distinguish SLE patients from healthy controls. (a) UMAP of latent factor matrix shows stratification between SLE patients and healthy controls (left). Violin plots (right) show the difference of factor values between SLE patients and healthy controls using Mann-Whitney U test. * adjust p-value < 0.05, **adjust p-value < 0.01. (b) UMAP of latent factor matrix shows uniform distribution of samples from different batches (left). Violin plots (right) shows the difference of factor values between batches using Mann-Whitney U test. * adjust p-value < 0.05, **adjust p-value < 0.01. (c) Scatter plots of factor 1 and 6 show stratification between SLE patients and healthy controls. (d) Heatmap with hierarchical clustering of factor 1 and 6 shows stratification between SLE patients and healthy controls. The annotation of columns (top) shows the difference of factor values between SLE patients and healthy controls using Mann-Whitney U test. * adjust p-value < 0.05, **adjust p-value < 0.01. (e) The heatmap displays percentage of variance explained for each factor across the different groups (batch) and views (cell type). (f) The heatmap displays correlation coefficient between different factors.

Up-regulation of interferon-related pathways shared across multiple cell types in SLE patients SLE related multicellular pathway modules consists of ‘Interferon alpha/beta signaling’ (type 1 interferon) in all cell types, ‘Type II interferon signaling (interferon-gamma)’ in nCM and classical dendritic cell (cDC), ‘Cross-presentation of particulate exogenous antigens (phagosomes)’ in cM and ‘Granzyme A-mediated apoptosis pathway’ in T8, etc. (Fig. 5a, Supplementary Table 2).

**Figure 5.**
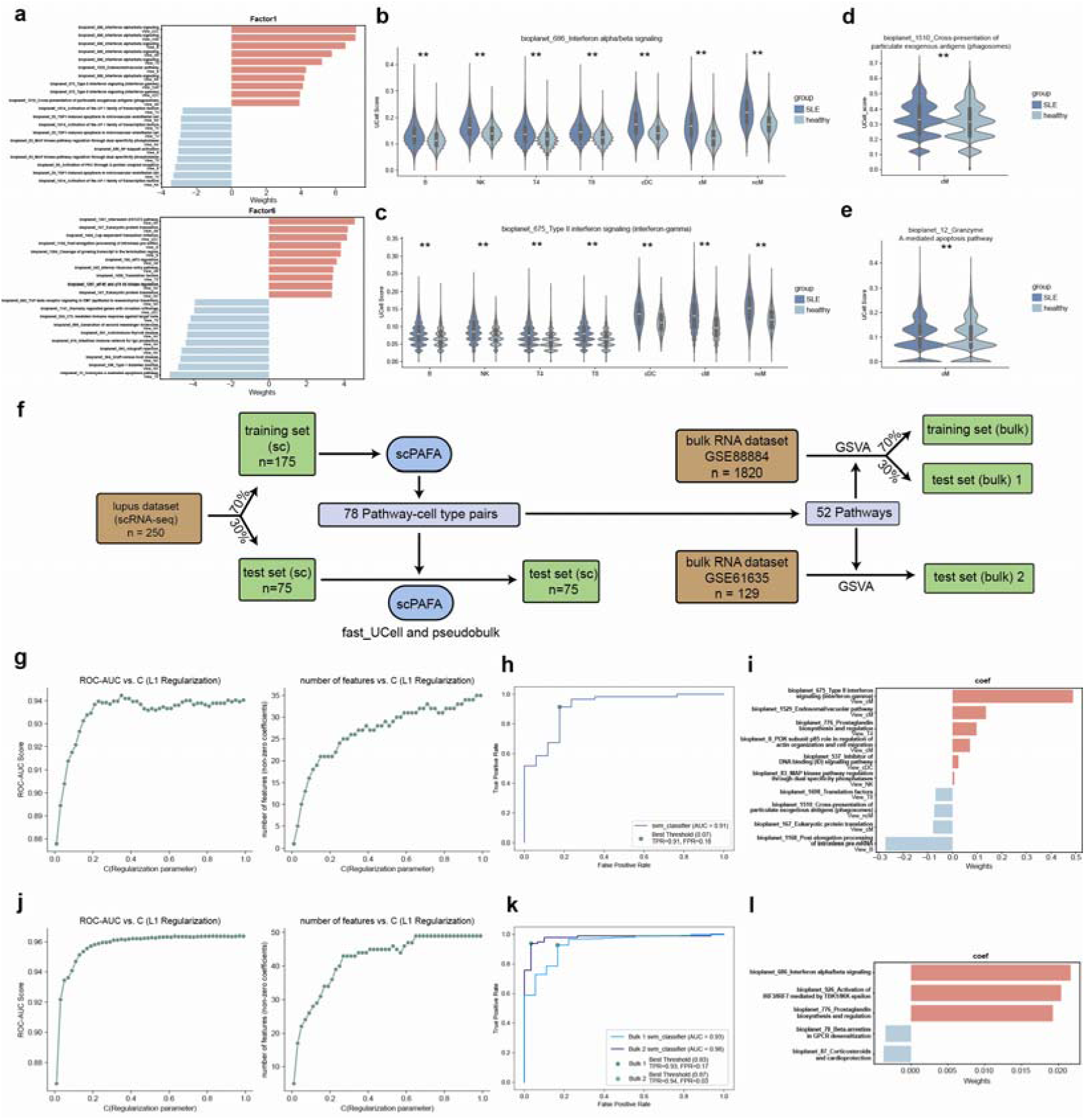
Pathway-centric interpretation and classifier training based on SLE related modules. (a) Butterfly bar plots displaying the pathway-cell type pairs with top 10 positive and negative weights of factor 1 and 6. (b) Violin plots show the difference of cell-level PAS of Interferon alpha/beta signaling’ between SLE patients and healthy controls using Mann-Whitney U test. * p-value < 0.05, ** p-value < 0.01. (c) Violin plots show the difference of cell-level PAS of ‘Type II interferon signaling (interferon-gamma)’ between SLE patients and healthy controls using Mann-Whitney U test. * p-value < 0.05, ** p-value < 0.01. (d) Violin plot shows the difference of cell-level PAS of ‘Cross-presentation of particulate exogenous antigens (phagosomes)’ between SLE patients and healthy controls using Mann-Whitney U test. * p-value < 0.05, ** p-value < 0.01. (e) Violin plot shows the difference of cell-level PAS of ‘Granzyme A-mediated apoptosis pathway’ between SLE patients and healthy controls using Mann-Whitney U test. * p-value < 0.05, ** p-value < 0.01. (f) Flowchart for classifier training on scRNA-seq data and bulk RNA data. (g) Line chart (left) illustrates the impact of various regularization parameter values on AUROC in the single cell training set (4-fold cross-validation). The line chart (right) illustrates the impact of different regularization parameter values on the number of features in the single cell training set (4-fold cross-validation). (h) The AUROC of the classifier on the single cell test set. (i) Butterfly bar plots displays the coefficients of features included in the classifier trained on single cell data. (j) Line chart (left) illustrates the impact of various regularization parameter values on AUROC in the bulk training set (4-fold cross-validation). The line chart (right) illustrates the impact of different regularization parameter values on the number of features in the bulk training set (4-fold cross-validation). (k) The AUROC of the classifier on the bulk test set and bulk independent test set. (l) Butterfly bar plots displays the coefficients of features included in the classifier trained on bulk data.

Conformingly, PAS of ‘ Interferon alpha/beta signaling’ and ‘ Type II interferon signaling (interferon-gamma)’ was significantly elevated across all cell types, especially in cDC and monocytes (cM,nCM) (Fig. 5b, c). Moreover, PAS of ‘Cross-presentation of particulate exogenous antigens (phagosomes)’ was significantly increased in cM (Fig. 5d), which could promote the activation of cytotoxic CD8+ T cells^36^, correspondingly, PAS of ‘Granzyme A-mediated apoptosis pathway’ was significantly upregulated in T8 of SLE patients (Fig. 5e). To further validate the reliability of the modules, next, we attempted to train SLE/healthy SVC classifier using SLE related multicellular pathway modules gained separately on scRNA-seq data and bulk RNA data. For scRNA-seq data, we used pseudobulk-PAS of 78 pathway-cell type pairs as features, which were the top-40 pathway-cell type pairs with the highest absolute weights associated with factors 1 and 6, respectively (Fig. 5f). After 4-fold cross-validation in training set, we set L1 regularization parameter as 0.05 for feature reduction (Fig. 5g), the AUROC of classifier achieved 0.91 in test set, which contained 10 pathway-cell type pairs (Fig. 5h, i). For bulk RNA data, 78 pathway-cell type pairs were aggregated into 52 pathways as features, 70% samples from GSE88884^37^ were assigned as bulk training set, while 30% samples from GSE88884 and all samples from GSE61635 were assigned as bulk test set 1 and 2 (Fig. 5f). Following training similar to abovementioned process, the AUROC of the 5-features-classifier including ‘Interferon alpha/beta signaling’ achieved 0.93 and 0.98 in bulk test set 1 and 2, separately (Fig. 5j-l).

## Discussion

In omics research, pathways are widely considered as informatics and functional biological units based on prior knowledge. Pathway analysis methods have been extensively employed with the objective of helping researchers to identify key biological themes for comprehending transcriptome^38, 39^. scPAFA can rapidly compute PAS for a large number of pathways on large-scale scRNA-seq data, allowing for efficient re-representation of transcriptome data based on prior knowledge. For PAS generating, we primarily used NCATS BioPlanet^26^ with scPAFA, a comprehensive integrated pathway resource that including manually-curated 1,658 publicly available human pathways, which have undergone comprehensive assessment for redundancy and consistency. Additionally, user can select appropriate pathway databases or use customized pathway set based on the specific biological context of the scRNA-seq dataset. For instance, we also employed 3CA metaprogram^27^ on CRC dataset for interpretation based on prior knowledge in oncology.

Cell type annotation is a key step in the analysis of scRNA-seq data, thus integrating transcriptomic features across different cell types to facilitate interpretation at the level of biological conditions is an issue worthy of attention. Several computational methodologies focused on multicellular integration have emerged: DIALOGUE^40^ was developed to systematically uncover combinations of coordinated cellular programs in different cell types from either spatial data or scRNA-seq data; Tensor-cell2cell^41^ can reveal context-dependent communication patterns linked to various phenotypic states, influenced by distinct combinations of cell types and ligand-receptor pairs; MOFAcellulaR^16^ allows the integration of measurements of independent single-cell, spatial, and bulk datasets to contextualize multicellular responses in disease. Inspired by the aforementioned methods, especially MOFAcellulaR^16^, we reutilized MOFA framework in scPAFA for multicellular integration. scPAFA can be regarded as a complement to existing pathway analysis methods in scRNA-seq data, aiming to identify interpretable pathway-based multicellular transcriptomic features associated with the biological conditions of interest.

In this study, scPAFA was applied to CRC and lupus dataset, identifying multicellular pathway modules that capture the transcriptomic features of MMR status in CRC and transcriptional abnormalities in SLE. These modules are consistent with previous reports in biological interpretation and can further be dissected to extract features for training classifiers. However, scPAFA also has a number of limitations. The selection of pathways is a supervised process, where using redundant pathway set may increase the complexity of the results, while choosing too few or pathways that do not align with the biological context may cause information loss. Moreover, pseudobulk integration of PAS matrix would lead to information loss since the PAS of multiple cells is aggregated. Besides, MOFA is a linear model, interpretability is achieved by sacrificing information content in each factor, the interpretability may also decrease with multiple disease states. In addition, the pathways and cell types within the same module can only suggest data-driven associations, rather than determining biological causal effects.

In summary, we present scPAFA, an efficient and user-friendly tool for researchers to generate PAS and uncover biologically interpretable disease-related multicellular pathway modules from scRNA-seq data. scPAFA is compatible with typical single-cell analysis workflows, which can integrate variance captured from different cell types and enhance understanding of disease biology, particularly in case-control cohort studies.

## METHODS

### Details of scPAFA framework

#### Input of scPAFA

The input for scPAFA includes AnnData class (built jointly with Scanpy) containing single-cell gene expression matrix and cell-level metadata, a python dictionary containing pathways name as keys and gene symbol in pathways as values, and a pandas dataframe containing sample-level metadata (clinical information). We provided a function ‘generate_pathway_input’ to filter and reformat pathway sets dictionary for subsequent analysis based on the number of overlapping genes between pathways and the gene expression matrix.

#### Details of ‘fast_ucell’

The function ‘fast_ucell’ is a python implementation of R package UCell^10^, which is faster, with similar adjustable parameters, capable of yielding consistent PAS. UCell calculates gene signature scores for scRNA-seq data based on Mann-Whitney U statistic, which is closely related to methods based on AUC scores such as AUCell^11^. UCell mainly consists of two steps: 1) Given single cell gene expression matrix *X* with genes as *g* and cells as *c* (e.g. count matrix or log-normalized count matrix), relative gene expression ranks matrix *R* is calculated by sorting all genes in descending order in each cell. To mitigate uninformative tail caused by the sparsity of single-cell data, *r_c,g_* = *r_max_* + 1 for all *r_c,g_* > *r_max_*, with *r_max_* = 1500 by default. 2) For a pathway *S* composed of genes (*S*_1_,…, *S_n_*), PAS of each cell j in *X* was calculated with the formula, ranging from 0 to 1:

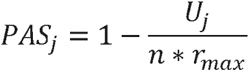

where *u_j_* is the Mann-Whitney U statistic calculated by:

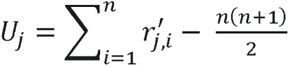

and 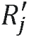 is obtained by sub-setting *R_j_* on the pathway *S*.

The performance of ‘fast_ucell’ running on large datasets was enhanced by multi-process parallelism and vectorized computation based on the original UCell computation process. In step 1, single cell gene expression matrix *X* in sparse matrix format from large dataset is automatically split into chunks of reduced size (default as 100,000 cells), the relative gene expression ranks matrix *R* is calculated parallelly by using ‘stats.rankdata’ function in scipy package, then *R* is reformatted into sparse matrix by assigning *r_c,g_* = 0 for all *r_c,g_* >,*r_max_* . In step 2, *R_sub_* is extracted from *R*, which consists of the union of genes contained in all pathways in pathway set *P*, then *R_sub_* is split into chunks by cells (default as 100,000 cells) for serially processing, while pathway set *P* is split into chunks by pathways for parallelly processing. For a pathway*S* composed of *n* genes, the PAS of *S* in a chunk of *R_sub_* is calculated by a vectorized computation process. In brief, *R*_S_ is extracted from the chunk of *R*_sub_, which only includes all genes in *S* ; row (cell)-wise summation of the *R*_S_ is performed and stored as vector *V*, and cells with a sum of 0 are labeled as *C*_0_, while others are labeled as *C*_1_. For cells in *C*_0_, the PAS of *S* is approximately specified as 0. For cells in *C*_l_, the numbers of zero columns(gene) in *R*_S_ are counted as vector *Z*, then PAS of *S* is calculated with the formula:

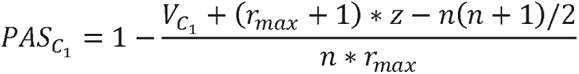

#### Details of ‘fast_score_genes’

The function ‘fast_score_genes’ is a multiprocessing implementation of ‘score_genes’ function in Scanpy (‘AddModuleScore’ function in Seurat). In function ‘score_genes’, PAS of a gene signature *S* in a given cell *j* of single cell gene expression matrix *X* is defined as the average relative expression of the overlap genes of *j* in *S*, which can be calculated by a three-step process: 1) using the average expression levels in all input cells to group all input genes into bins (default as 25 bins) of equal size, 2) randomly selecting reference genes (default as 50 genes) from the same expression bin for each gene in the gene signature *S* as reference gene set *S_ref_* 3) the PAS of cell *j* is the average expression of *S* subtracted with the average expression of reference gene set *S_ref_* :

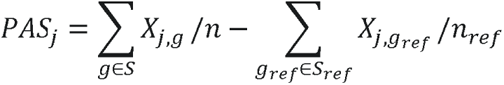

where *X_j,g_* is the expression of gene *g* in cell *j*, *n* and *n_ref_* is the number of genes in *S* and *S_ref_*, respectively. In ‘fast_score_genes’, given a single cell gene expression matrix *X*, step 1 was performed first, then *X* can be split into smaller chunks by cells (default as 100,000 cells) for serially processing (minimizing memory usage). For each chunk of *X*, steps 2 and 3 can be executed concurrently using multiple pathways in pathway set *P*.

#### Details of pseudobulk integration functions

We developed ‘generate_scpafa_input’ and ‘generate_scpafa_input_multigroup’ functions for pseudobulk integration with PAS matrix and cell-level metadata. Users can specify the column names in cell-level metadata corresponding to sample information, cell type, and batch information as well as quality control metrics such as the cell number threshold of a pseudobulk sample, the sample number threshold and the sample proportion threshold within groups of a cell type. These functions generate a long-format table, which is the input for MOFA containing 5 columns: column ‘sample’ including sample information, column ‘group’ including batch information, column ‘feature’ including names of pathways, column ‘view’ including cell type information and column ‘value’ including pseudobulk PAS values.

#### Wrapper function for MOFA model training

We integrated the source code from mofapy2 (v0.7.0) with modifications to support the new version of the pandas (≥v2.0) package. For single-click runnable MOFA model training, we provided wrapper function ‘run_mofapy2’ based on the ‘entry_point’ class of mofapy2.

#### Downstream analysis functions

Python package mofax (v0.3.6) is employed for extracting latent factor matrix and weight matrix from the converged MOFA model. For downstream analysis, we provided ‘parametric_test_category’ and ‘nonparametric_test_category’ functions to identify disease-related multicellular pathway modules from latent factor matrix and sample-level metadata, besides, we also provided ‘cal_correlation’ function to identify modules associated with continuous traits. For visualization, ‘runumap_and_plot’, ‘plot_factor_scatter_2D’ and ‘draw_cluster_heatmap’ functions are used to show stratification of samples, while ‘plot_weights_butterfly’ function can be used to show features with top absolute weights.

### Single-cell RNA-seq data preprocessing

For CRC dataset^22^, single cell gene expression matrix and metadata was downloaded from Broad Institute’s Single Cell Portal (https://singlecell.broadinstitute.org/single_cell/study/SCP1162). We used Scanpy (v1.9.5) to bulid AnnData, only genes with gene ID starting with ’ENSG’ were retained, then after filtering out genes expressed in fewer than 30 cells, the dataset comprises 371,223 cells and 26320 genes. For lupus dataset^3^, AnnData in h5ad format was downloaded from CZ CELLxGENE Annotate(https://cellxgene.cziscience.com/collections/436154da-bcf1-4130-9c8b-120ff9a888f2), after filtering out genes expressed in fewer than 30 cells, the dataset includes 1,263,676 cells and 20514 genes.

### Bulk RNA data preprocessing

Microarray gene expression matrix and corresponding sample-level metadata of GSE39582, GSE61635 and GSE88884 was collected using GEOparse(v2.0.3). Bulk RNA-seq gene expression matrix and corresponding sample-level metadata of TCGA Colon and Rectal Cancer (COAD-READ) dataset was collected from UCSC Xena(https://xena.ucsc.edu/), samples from primary solid tumor were used for further analysis. After filtering out samples with missing clinical information, TCGA COAD-READ dataset comprises 251 MMRp samples and 53 MMRd samples; GSE39582 includes 444 MMRp samples and 72 MMRd samples; GSE61635 contains 30 healthy controls and 99 SLE samples, while GSE88884 includes 60 healthy controls and 1760 SLE samples.

### Runtime and memory usage evaluation

We used an Intel X79 server (Ubuntu 20.04) with E5-2670v2 cpu and 256GB RAM for runtime and memory usage evaluation. For CRC dataset, we sampled 100 thousand, 200 thousand, and all cells from the dataset as input; the pathways from NCATS BioPlanet collection and 3CA metaprogram with more than 6 overlap genes (1629 pathways) were used as pathway input. For lupus dataset, we sampled 100 thousand, 200 thousand, 500 thousand, and all cells from the dataset as input, the pathways from NCATS BioPlanet collection with more than 6 overlap genes (1383 pathways) were used as pathway input. UCell (v2.6.2) and ‘score_genes’ from Scanpy (v1.9.5) was used for comparing. To ensure consistent computational performance each time the program is executed, we conduct all tests sequentially.

### Application of scPAFA on CRC single cell dataset

We extracted 108,497 tumor epithelial cells from the CRC dataset AnnData as the input for scPAFA. Python dictionary containing NCATS BioPlanet with 1,658 pathways and 3CA metaprogram with 149 gene sets was used as pathway input. After filtering out pathways with fewer than 6 overlap genes with the expression matrix, 1629 pathways remained. PAS matrix was calculated using ‘fast_ucell’ function, setting max rank (*r_max_*) as 2000. Firstly, we used single-group MOFA framework to identify potential batch effects, function ‘generate_scpafa_input’ was used for PAS pseudobulk integration with 65 pseudobulk samples (30 MMRp and 35 MMRd), specifying cell type as ‘view’, setting a pseudobulk sample to include at least 10 cells, and a ‘view’ to consist of at least 45 pseudobulk samples. After identifying batch effect in samples’ stratification caused by different 10X chemistry version, we used ‘generate_scpafa_input_multigroup’ to aggregate PAS matrix for multi-group MOFA+ framework, specifying batch as ‘group’, setting a ‘view’ to consist of more than 75% of pseudobulk samples (at least 15) in each group. In MOFA model training, we set factor number as 10 and filtered out factors explained less than 1% of variation. Latent factor matrix and weight matrix from the converged MOFA model was extracted using mofax. We employed ‘nonparametric_test_category’ functions with latent factor matrix and sample-level metadata to identify MMR status related multicellular pathway modules by significance of Mann-Whitney U test. The aforementioned visualization functions are utilized to illustrate stratification of samples and biological interpretation of modules.

### Application of scPAFA on lupus single cell dataset

The lupus dataset is based on multiplex scRNA-seq with cell-level processing batch information^3^. We extracted 1,067,499 cells from 3 processing batches that simultaneously contain SLE and health controls as scPAFA input, along with 1383 pathways from NCATS BioPlanet which contain more than 6 overlap gene with the expression matrix. We generated PAS matrix by using ‘fast_ucell’ function, setting max rank (*r_max_*) as 2000. Subsequently, we employed single-group MOFA framework to validate batch effect caused by processing batch; for corresponding pseudobulk integration, we set a pseudobulk sample to include at least 20 cells, and a ‘view’ to consist of at least 220 pseudobulk samples. We observed batch effect in samples’ stratification caused by processing batch, we also identified a cluster of 32 pseudobulk samples that may be contaminated by red blood cells by using k-means clustering and scPAFA downstream analysis functions. After filtering out this cluster of samples, the remained 250 pseudobulk samples were split into a training set with 175 pseudobulk samples (51 healthy controls and 124 SLE) and a testing set with 75 pseudobulk samples (17 healthy controls and 58 SLE). The PAS matrix with 664,214 cells from training set was aggregated then used for multi-group MOFA+ framework, specifying processing batch as ‘group’, setting a ‘view’ to consist of more than 75% of pseudobulk samples (at least 15) in each group. Similar to the steps described in CRC dataset, MOFA model was trained and SLE related multicellular pathway modules were identified and interpreted.

### Machine learning model training and evaluation

For CRC dataset, top-30 pathway-cell type pairs with the highest absolute weights associated with factors 2, 3 and 6 were collected. After removing cell type information and eliminating duplicate values, the PAS of the remained 30 pathways were calculated on GSE39582 and TCGA COAD-READ data using ‘gsva’ function in gseapy (v1.1.0) and used as features for classifier training. GSE39582 was split into a 70% training set and a 30% test set, whereas TCGA COAD-READ data was used as independent test set. We used ‘LinearSVC’ function with L1 regularization from scikit-learn (v 1.3.2) to train MMR status classifier. After determine the parameter for L1 regularization by employing 4-fold cross-validation on the training set, we set C as 0.2 and trained SVC model. The AUROC of converged model on test set and independent test set was characterized to evaluate the performance of the model.

For lupus dataset, SLE/healthy SVC classifier was trained on scRNA-seq data and bulk RNA data, separately. For scRNA-seq data, PAS of 78 pathway-cell type pairs were extracted from pseudobulk PAS matrix and used as features for classifier training, which were collected and eliminated duplicate from top-40 pathway-cell type pairs with the highest absolute weights associated with factors 1 and 6. After determine the parameter for L1 regularization by employing 4-fold cross-validation on the training set with 175 pseudobulk samples, we set C as 0.05 and trained SVC model. The AUROC of the classifier was evaluated on the test set with 75 pseudobulk samples.For bulk RNA data, 78 pathway-cell type pairs were aggregated into 52 pathways then used as features for classifier training. SLE/healthy SVC classifier was trained and evaluated following similar process in abovementioned CRC dataset, while using 70% of GSE88884 as training set, 30% of GSE88884 as test set, GSE61635 as independent test set and L1 regularization parameter C as 0.01.

### Statistical information

In our study, the difference of latent factor values between different disease states was examined by Mann-Whitney U test and Benjamini-Hochberg correction for p-values was applied. The violin plots showing the difference of cell-level PAS between different disease states were also examined by Mann-Whitney U test. In the figures, p-values or adjusted p-values exceeding 0.05 were deemed statistically non-significant and left unlabeled. Values equal to or below 0.05 and 0.01 were denoted with * and **, respectively.

### Data availability

CRC scRNA-seq data can be accessed at Broad Institute’s Single Cell Portal (https://singlecell.broadinstitute.org/single_cell/study/SCP1162). Lupus scRNA-seq data can be accessed at CZ CELLxGENE Annotate(https://cellxgene.cziscience.com/collections/436154da-bcf1-4130-9c8b-120ff9a888f2). GSE39582, GSE61635 and GSE88884 is publicly accessible. TCGA COAD-READ RNA-seq data can be found via UCSC Xena browser(https://xena.ucsc.edu/).

### Code availability

scPAFA is open-source and publicly hosted on GitHub (https://github.com/ZhuoliHuang/scPAFA) with documentation and tutorials.

## Supporting information

Supplementary Table 1. Weights matrix of CRC dataset, related to Figure 3a.

Supplementary Table 2. Weights matrix of lupus dataset, related to Figure 5a.

## Acknowledgments

We acknowledge China National GeneBank (CNGB) for its support for this study.

## Author contributions

J.Y, and X.J. led and supervised the study. Z.H. designed and implemented the scPAFA software, and led the data analyses. Z.H., Y.Z. and W.W. created visualization of result. J.Y, W.Z., and C.W. provided biological interpretation. Z.H., J.Y, X.Z., and X.J. wrote the manuscript.

---

**Supplementary Figure 1.**
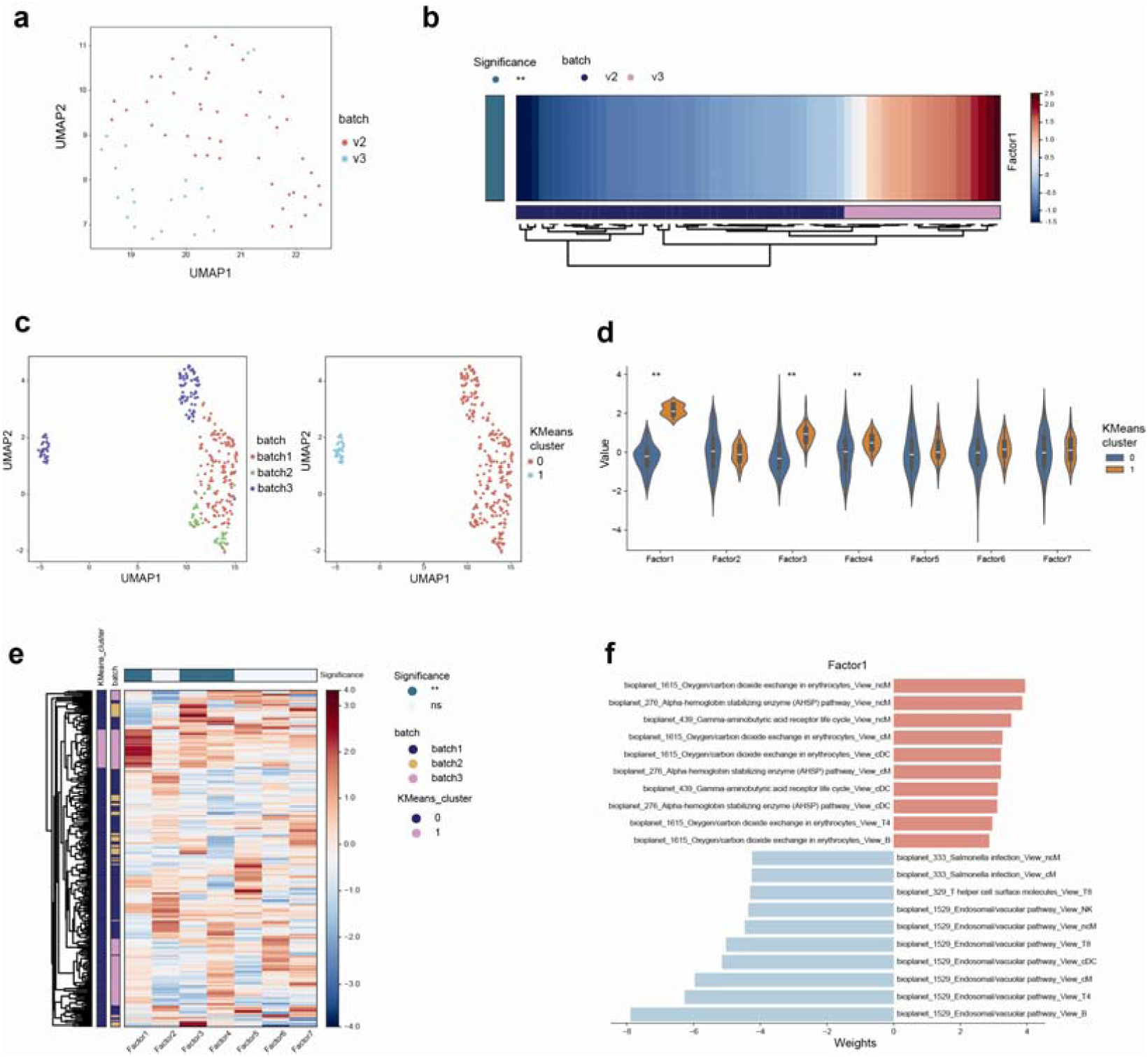
Batch effects captured by single-group MOFA framework. (a) UMAP of latent factor matrix shows stratification in CRC dataset caused by batch. (b) Heatmap of factor 1 shows stratification in CRC dataset caused by batch. The difference of factor 1 values between MMRp and MMRd samples was examined using Mann-Whitney U test. * adjust p-value < 0.05, **adjust p-value < 0.01. (c) UMAP of latent factor matrix shows stratification in lupus dataset caused by processing batch (left). Two distinct k-means clusters on UMAP plot (right). (d) Violin plots (right) shows the difference of factor values between k-means cluster 0 and 1 using Mann-Whitney U test. * adjust p-value < 0.05, **adjust p-value < 0.01. (e) Heatmap with hierarchical clustering of latent factor matrix shows stratification between k-means cluster 0 and 1. The annotation of columns (top) show the difference of factor values between k-means cluster 0 and 1 using Mann-Whitney U test. * adjust p-value < 0.05, **adjust p-value < 0.01. (f) Butterfly bar plots displaying the pathway-cell type pairs with top 10 positive and negative weights of factor

